# Exploring the genetic diversity of the Japanese Population: Insights from a Large-Scale Whole Genome Sequencing Analysis

**DOI:** 10.1101/2023.01.23.525133

**Authors:** Yosuke Kawai, Yusuke Watanabe, Yosuke Omae, Reiko Miyahara, Seik-Soon Khor, Eisei Noiri, Koji Kitajima, Hideyuki Shimanuki, Hiroyuki Gatanaga, Kenichiro Hata, Kotaro Hattori, Aritoshi Iida, Hatsue Ishibashi-Ueda, Tadashi Kaname, Tatsuya Kanto, Ryo Matsumura, Kengo Miyo, Michio Noguchi, Kouichi Ozaki, Masaya Sugiyama, Ayako Takahashi, Haruhiko Tokuda, Tsutomu Tomita, Akihiro Umezawa, Hiroshi Watanabe, Sumiko Yoshida, Yu-ichi Goto, Yutaka Maruoka, Yoichi Matsubara, Shumpei Niida, Masashi Mizokami, Katsushi Tokunaga

**Author notes:** Corresponding authors (Y.K.); (K.T.). Department of Biological Sciences, Graduate School of Science, The University of Tokyo, Bunkyo-ku, Tokyo 113-8654, Japan (Yusuke Watanabe). Center for Surveillance, Immunization and Epidemiologic Research, National Institute of Infectious Diseases, Shinjuku-ku, Tokyo 162-8640, Japan (Reiko Miyahara).

## Abstract

The Japanese archipelago is a terminal location for human migration, and the contemporary Japanese people represent a unique population whose genomic diversity has been shaped by multiple migrations from Eurasia. Through high-coverage whole-genome sequencing (WGS) analysis of 9,850 samples from the National Center Biobank Network, we analyzed the genomic characteristics that define the genetic makeup of the modern Japanese population from a population genetics perspective. The dataset comprised populations from the Ryukyu Islands and other parts of the Japanese archipelago (Hondo). Low frequency detrimental or pathogenic variants were found in these populations. The Hondo population underwent two episodes of population decline during the Jomon period, corresponding to the Late Neolithic, and the Edo period, corresponding to the Early Modern era, while the Ryukyu population experienced a population decline during the shell midden period of the Late Neolithic in this region. Genes related to alcohol and lipid metabolism were affected by positive natural selection. Two genes related to alcohol metabolism were found to be 12,500 years out of phase with the time when they began to be affected by positive natural selection; this finding indicates that the genomic diversity of Japanese people has been shaped by events closely related to agriculture and food production.

**Author summary:** The human population in the Japanese archipelago exhibits significant genetic diversity, with the Ryukyu Islands and other parts of the archipelago (Hondo) having undergone distinct evolutionary paths that have contributed to the genetic divergence of the populations in each region. In this study, whole genome sequencing of healthy individuals from national research hospital biobanks was utilized to investigate the genetic diversity of the Japanese population. Haplotypes were inferred from the genomic data, and a thorough population genetic analysis was conducted. The results indicated not only genetic differentiation between Hondo and the Ryukyu Islands, but also marked differences in past population size. In addition, gene genealogies were inferred from the haplotypes, and the patterns were scrutinized for evidence of natural selection. This analysis revealed unique traces of natural selection in East Asian populations, many of which were believed to be linked to dietary changes brought about by agriculture and food production.

## Introduction

The Japanese archipelago is located in the eastern part of the Eurasian continent and is one of the final destinations of the human migration out of Africa. While the identity of the first human groups to reach the Japanese archipelago is uncertain, the Jomon people, who were hunter-gatherers known for their pottery, lived in the region after 16,000 years ago. The genetic diversity of the peoples of the Japanese archipelago underwent a dramatic transformation following the Yayoi Period, which began around 3,000 years ago with the migration of agriculturalists from Eurasia. Genome analysis of ancient and modern humans has shown that they admixed with the originally inhabited Jomon people, resulting in the genetic diversity of the modern Japanese population from the Yayoi people. This process is thought to have started on Kyushu Island and then gradually spread throughout the archipelago. Although the Ryukyu Islands are separated from Kyushu Island by a significant distance, the agricultural culture known as the Gusuku period began 800 years ago, and it is believed that this agriculture was introduced by migrants from the mainland. These long histories of migration and admixture have shaped the genetic diversity of the peoples of the Japanese archipelago. Previous genome analyses have attempted to reveal this genetic diversity, but most have only sampled specific regions and therefore have been insufficient to fully examine the genetic diversity of the Japanese archipelago as a whole. In this study, we used whole-genome sequencing data from subjects from a wide range of regions in the Japanese archipelago to more fully understand the genetic diversity of the peoples of the archipelago.

There are six national research hospitals in Japan that specialize in advanced medical care and research, and each of them maintains its own biobank that collects and stores biological samples from patients. National Center Biobank Network (NCBN) is a federation of these centers that collaborate to provide samples, genomic and clinical information, and public relations. In this study, we performed WGS of 9,850 individual DNA specimens stored in the biobanks of NCBN. These biobanks are located in three distinct regions of Japan: NCGM, NCCHD, and NCNP in the Tokyo area; NCGG in Aichi Prefecture in the central area in the Honshu Island; and NCVC in Osaka Prefecture in the western area in Honshu Island. Therefore, the genomic information obtained in the present study is expected to reflect the regional diversity of Japan to some extent. Here, we characterized the data obtained from this analysis and described the genetic diversity in the Japanese population.

## Results

### Whole genome sequencing analysis

DNA samples from 9,850 individuals from five National Center Biobanks were analyzed using WGS, and the data in FASTQ format were received from the outsourced laboratory. The received data were processed through the primary data analysis pipeline to obtain mapping results and variant call results. Quality control (QC) metrices were calculated to confirm that stable quality analysis was obtained (Fig 1). The autosomes had an average read depth of 34.0 ± 2.4, and the average insert length of the reads was 703 +/- 30 bp. The mapping rate per sample was 99.99% ± 0.39%. These statistics did not vary significantly between samples, and there was no clear bias between biobanks except for saliva samples. The saliva samples showed lower mapping rates than the blood samples, probably due to the foreign DNA in the saliva.

**Fig 1.**
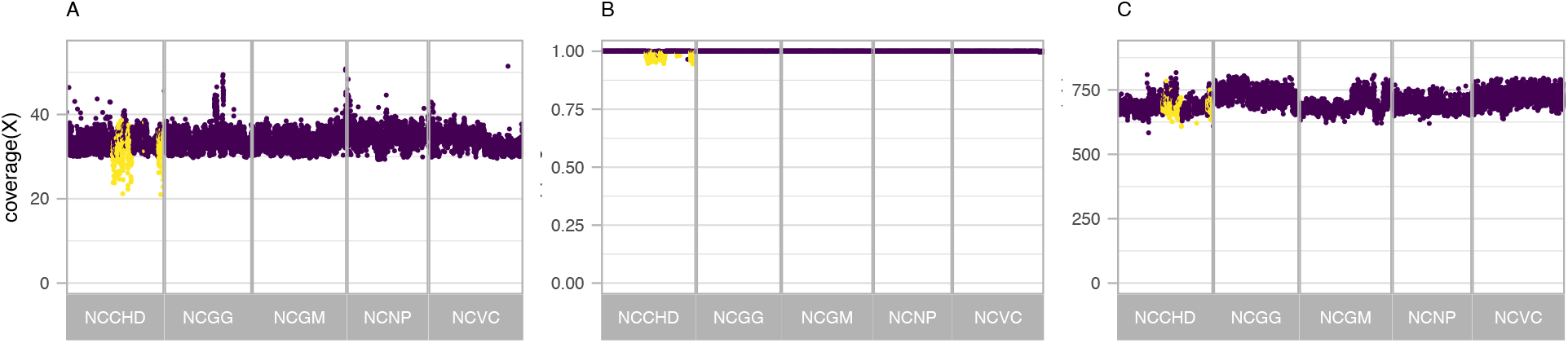
Quality control metrics of whole-genome sequencing. Quality control metrics for each sample are plotted against the sample in the horizontal axis direction; QC indices are (A) average coverage of reads in autosomal loci after excluding duplicated reads, (B) mapping rate, and (C) average insert length. Saliva-derived samples are colored by yellow.

### Summary and accuracy of SNP and short insertion and deletions

Variants were characterized by joint calling to integrate individual variant information. In this analysis, we performed joint calling of the gVCFs of 9,287 samples from NCBN, analyzed at the time of writing, together with 2,504 samples from the International 1000 Genomes project (S1 Table). The VCF obtained after the joint call contained a total of 208,785,859 records, of which 88.5% (184,864,563) passed the filtering using Variant Quality Score Recalibration (VQSR). We found 122,459,307 variants after focusing only on the variants in the NCBN samples, and 86.3% (105,729,588) of them passed the filter of VQSR. Of the variants that passed the filter, 87,246,166 were single nucleotide variants (SNVs) and 18,483,422 were short insertion and deletions (INDELs); 47% (41,046,547) of the SNVs and 39.8% (7,361,318) of the INDELs that passed the filter were novel variants not registered in dbSNP151. Most of the novel variants were very rare. For example, 34.56% of the known SNVs were singletons found as the heterozygous genotypes of one sample out of 9,287 individuals, and 86.73% were observed at a very low frequency of less than 0.5%. Conversely, 67.46% of the novel variants not registered in dbSNP were singletons, and the percentage of SNVs with a frequency less than 0.5% was more than 99.99%. This is consistent with previous reports that most novel variants are found privately [3–6].

We evaluated the accuracy of the variants using two approaches. First, we performed the genotyping using SNP array to estimate the degree of genotype concordance with WGS results. For this purpose, genome-wide genotyping using the SNP array on the DNA samples of 448 individuals who had undergone WGS analysis was conducted. The 639,508 autosomal SNPs remaining after variant QC in the SNP array were compared with the results obtained after WGS analysis. The number of mismatches ranged from 66 to 7,205 per sample, with an average of 408.7. As a result, the average discordance rate between the two sets of variants was 0.063%. This estimate appears to be a conservative estimate of the error, as it is primarily concentrated in a region that is easy to analyze and for which probes are designed on SNP arrays. Then, we compared the genotypes of the trio samples to estimate the frequency with which the offspring of a trio had heterozygous or non-reference homozygous variants whose parents’ genotypes did not follow the pattern expected from Mendelian low. The sample analyzed in this study contained 148 trios of parents and offspring. We observed the inheritance pattern of genotypes from an average of 4,284,264 variants per trio. Of these, 6,448.4 (0.15%) had an abnormal inheritance. This percentage became more pronounced when stratified by the novelty of the variants, e.g., the known SNVs and INDELs had error rates of 0.09% and 0.42%, respectively, with errors in the inheritance pattern, whereas the novel SNVs and INDELs had error rates of 2.26% and 10.9%, respectively. The Mendelian heritability errors can include the sequencing or genotyping error, *de novo* mutations, and gene conversions in the parents’ gametes. However, our estimates approximate the error rate of genotyping in this study.

### Ancestry inference and allele frequency distribution

We conducted the principal component analysis (PCA) to identify the ancestry of NCBN samples. After removing 20 samples with a call rate below 95%, Identical-by-Descent (IBD) was used to detect related samples, resulting in 8,972 and 2,493 unrelated samples from NCBN and the International 1000 Genomes project, respectively. PCA using these samples detected 21 NCBN samples not belonging to the East Asian populations (Fig 2A). Furthermore, when PCA was performed only on East Asians, the samples were divided into two clusters: one consisting of continental populations (Han Chinese in Beijing; CHB, Han Chinese South; CHS, Kinh Vietnamese; KHV, Chinese Dai in Xishuangbanna, China; CDX) and the other including Japanese in Tokyo (JPT) from 1000 Genomes and NCBN samples (Fig 2B). In addition, the latter cluster was divided into large and small clusters consistent with the previous studies [7–9] in which the larger one was called the Hondo population and the smaller one was called the Ryukyu population [7]. In this study, we followed this convention (S1 Fig). The Hondo cluster consisted of 8,524 people, whereas the Ryukyu cluster consisted of 182 people. We compared the allele frequencies of the Japanese population (GEM Japan Whole-genome Aggregation) estimated based on the WGS analysis in previous studies with those of the Hondo sample and found significant frequency agreement (Fig 3A). While the allele frequencies between the Hondo and Ryukyu populations also showed high agreement, the breadth of the distribution was wider than the comparison between Hondo and GEM Japan (Fig 3B). This could be due to the difference in the mainland and Ryukyu populations and the subsequent genetic drift.

**Fig 2.**
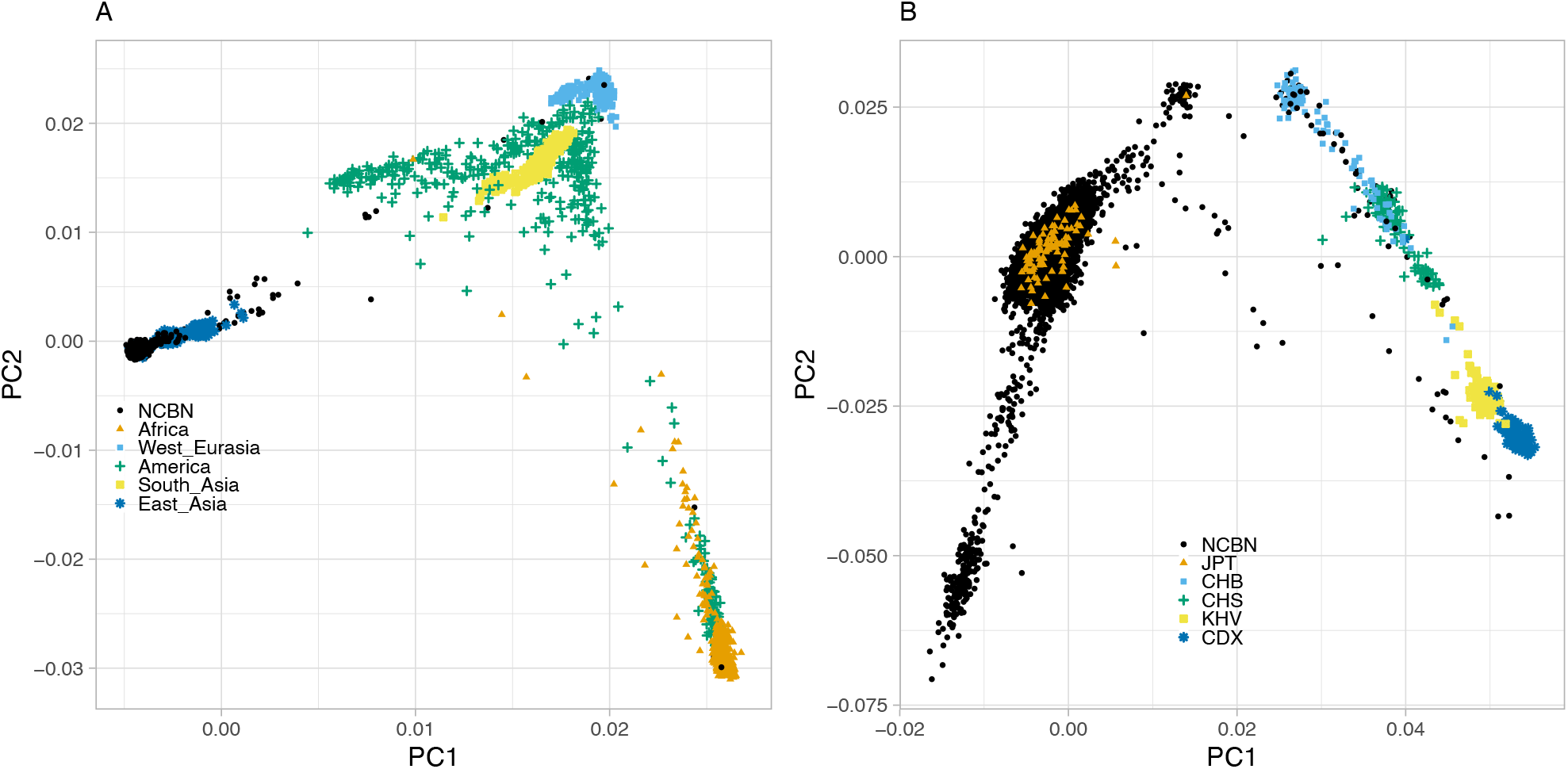
Genetic structure of NCBN samples. (A) The first and second principal components are plotted. The continental population of the international 1000 genomes and NCBN are plotted in different colors and shapes. (B) PCA plots of the East Asian population of the International 1000 Genomes and NCBN samples are shown. JPT: Japanese in Tokyo, Japan, CHB: Han Chinese in Beijing, China, CHS: Han Chinese South, KHV: Kinh in Ho Chi Minh City, Vietnam, CDX: Chinese Dai in Xishuangbanna, China

**Fig 3.**
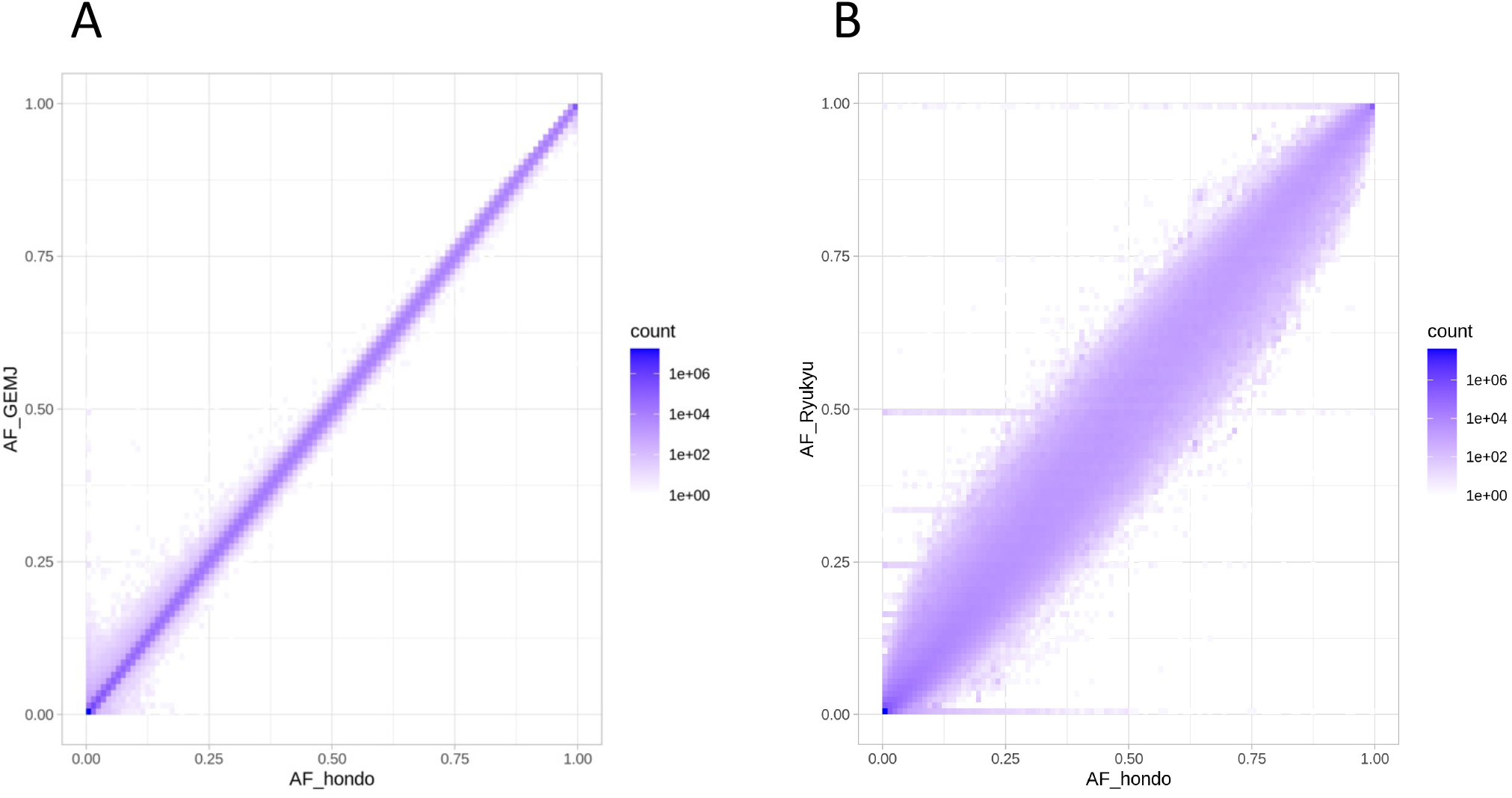
Comparison of allele frequency between different populations. (A) The non- reference allele frequencies of the Hondo population of NCBN samples (X-axis) and the corresponding variants of GEM Japan (Y-axis) were counted and then the numbers were plotted as density. (B) Same plot for Hondo population (X-axis) and Ryukyu population (Y-axis).

### Functional landscape of variants

The variants identified by WGS analysis were annotated for their biological functions. The impact of the variants was classified based on the criteria of the annotation software and the database as described in the Methods section. Deleterious mutations are more likely to be kept at low frequencies in the population, as such mutations are less likely to spread in the population because of negative selection. In fact, variants with a high impact on annotation showed a clear tendency to have a low frequency in the population. The LOFTEE plugin of Variant Effect Predictor was used to detect loss-of-function (LoF) variants in the Hondo and Ryukyu populations. For comparison, we also detected LoF variants in 26 populations in the 1000 Genomes Project phase 3 dataset [10]. 14,145 SNVs and 16,823 INDELs were detected as high confident LoF specific to the Hondo population. For the Ryukyu population, 211 SNVs and 288 INDELs were detected. The vast majority of LoF SNVs exhibited a very low frequency in the Hondo population (Fig 4A). In fact, 76.0% of these SNVs exhibited allele frequencies below 0.01%. We compared the number of LoF alleles and the number of homozygous of LoF alleles per individual for Hondo, Ryukyu, and populations in the 1000 Genomes Project (Fig 4 B and C, S2 Fig). Since homozygous LoFs result in a complete loss of gene function, the number of homozygous LoFs in an individual’s genome can be used to measure the individual’s genetic burden. Both indices were highest in Africa, lowest in West Eurasia, and moderate in Hondo and Ryukyu. The number of homozygous LoF alleles per individual by allele frequency was generally higher in Africa across all allele frequencies (Fig 4D), which is consistent with the trend observed in a previous study [11].

**Fig 4.**
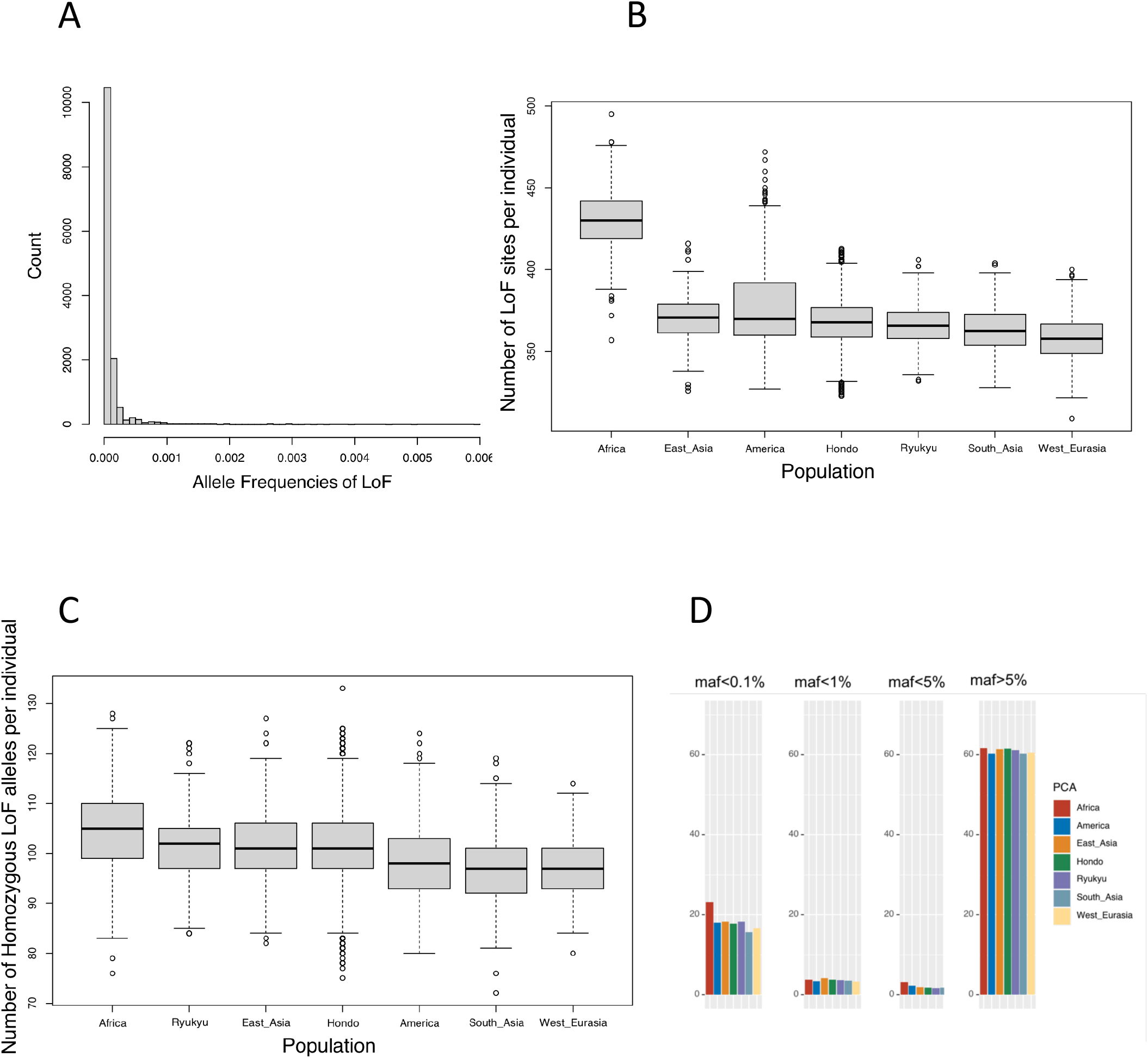
Analysis of loss-of-function (LoF) variants. (A) The allele frequency distribution of newly detected HC LoF SNPs in the Hondo population. (B) The number of LoF alleles and (C) the number of homozygous of LoF alleles per individual for Hondo Ryukyu, and populations of the International 1000 Genomes. (D) The number of homozygous of LoF alleles per individual by allele frequency for Hondo, Ryukyu, and the populations of the International 1000 Genomes.

We compared the variants of NCBN samples with ClinVar registered variants [12]. A total of 103,833 variants found in the Hondo population are registered in ClinVar. Of these, 2,427 were classified as “pathogenic” or “likely pathogenic” variants. Seven variants were found in the four-star category, the most reliable classification based on the ClinVar review status. Only one of them was “pathogenic” and a singleton variant (i.e., heterozygous in a person) of the CTFR gene. The remaining six were polymorphic variants related to drug responsiveness of CYP2C19. There were 1,130 variants in the 3-star category reviewed by the expert panel. Of these, 56 were “pathogenic,” and 13 were “likely pathogenic.” The frequencies of these variants were the highest, at 1.0%, and most of them were extremely rare; only a few were observed in the population. Most of the less well-reviewed variants with <3 stars had frequencies of less than 1%, but 34 variants had a frequency of 1% or more.

### Allele frequency estimation of HLA loci

Three-field HLA calling results from the WGS dataset in the present study were compared with HLA allele frequencies HLA Foundation Laboratory (Kyoto, Japan) (S3 Fig). All common HLA alleles (frequencies >1%) were concordant between the two datasets with observed differences of less than 1%. To further validate our HLA calling results, a subset of 94 samples was subjected to high-resolution HLA genotyping. Three-field HLA class I (HLA-A, -C, and -B) accuracies were 96.3%, 97.9%, and 96.8%, respectively, and 3-field HLA class II (HLA-DRB1, -DQA1, -DPA1, and -DPB1) accuracies were 98.9%, 100.0%, 98.9%, 100.0%, and 96.8%, respectively. The accuracy of 2-field HLA class I (HLA-A, -C, and -B) increased to 97.9%, 98.4%, and 97.3%, respectively.

### Evolutionary perspective of genomic diversity

The recent decrease in population size was detected in Hondo and Ryukyu populations. Figure 5A shows the population histories of Hondo and Ryukyu populations inferred using IBDNe [13], which estimated the changes in population size in recent (∼200 generations ago) past based on IBD sharing among individuals. In Hondo, the population size decreased from about 75 to 50 generations ago, and from 17 to 11 generations ago. In the Ryukyu population, a reduction in population size was observed from about 100 to 25 generations ago. The distributions of IBD length were multimodal in both populations, indicating fluctuations in population size (Fig 5B and 5C). We also estimated the long-term changes in the effective population size from the genome-wide genealogy using Relate software [14]. We estimated genome-wide genealogy based on the whole-genome data of 1,000 randomly selected samples from Hondo, 182 samples from the Ryukyu, and the CHB population from the 1000 Genomes Project. The Ryukyu population showed a bottleneck that peaked around 2,700 years ago (S4 Fig). Hondo/CHB population and Ryukyu population diverged around 3,700 years ago, consistent with previous estimations of the divergence time using SNP arrays [15, 16].

**Fig 5.**
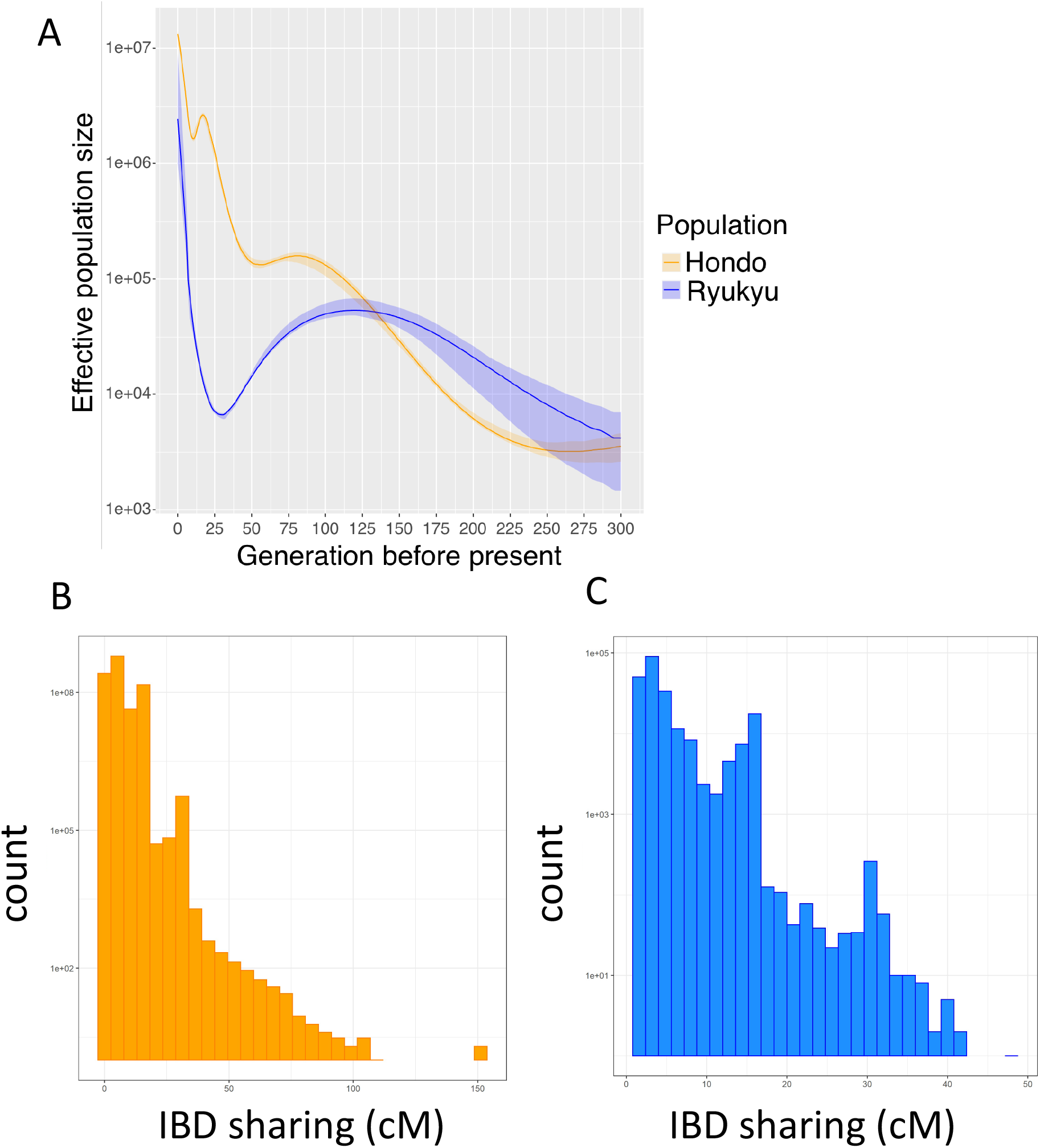
Estimation of past population size from IBD sharing. (A) Short-term effective population size change in Hondo and Ryukyu populations by IBDNe. (B) Distribution of IBD segment length in Hondo. (C) Distribution of IBD segment length in Ryukyu.

We detected positive natural selection based on genome-wide genealogy of 1,000 Hondo samples and found SNPs with p-values below the genome-wide significance level (p < 5.0 × 10^-8^) (S5 Fig, S2 Table). As the QQ plot suggested inflation of the test statistics (S6 Fig), it is possible that the results contain false positives. However, the genes reported in previous studies, which may have undergone positive natural selection, were correctly included in the results. It is therefore important to consider this when interpreting the results. For example, ALDH2 rs671 G/A (p-value = 2.0 × 10^-17^) and ADH1B rs1229984 T/C (p-value = 6.8 × 10^-10^), which are associated with alcohol metabolism, showed positive natural selection signals [17, 18]. The genealogies of the genes showed that the derived alleles were spreading rapidly through the population (S7 Fig). The second example is signals of positive natural selection on the non-synonymous rs76930569 C/T (p-value = 1.1 × 10^-12^) variant in the OCA2 gene. This variant is in complete linkage equilibrium with rs1800414 T/C, involved in melanin biosynthesis, and has been shown to be associated with light skin color and tanning ability in Asian populations [19–21]. The third example of the positive selection is the FADS gene family. Multiple SNPs (rs174599, rs174600, rs174601, rs97384, rs57535397, rs76996928) showed the signatures of positive selection. FADS1 and FADS2 encode catalytic proteins, which synthesize long-chain fatty acids from short-chain fatty acids [22], and have been subjected to natural selection related to diet in several human populations [22–26]. We further analyzed change in allele frequency with time for these genes under positive natural selection. We used CLUES software [27] to estimate the allele frequency trajectory of SNPs in ALDH2, ADH1B, OCA2, and the FADS gene family. The frequency of the derived alleles in ADH1B rs1229984 increased about 20,000 years ago (Fig 6A). In contrast, the frequency of ALDH2 rs671 increased from about 7,500 years ago (Fig 6B). The allele frequency trajectory of OCA2 rs1800414 (Fig 6C) showed that the frequency of derived allele of OCA2 rs1800414 began to increase due to natural selection around 25,000 years ago. The frequency of derived allele of rs174599 began increasing around 25,000 years ago, slowed down 15,000 years ago, and started increasing again 10,000 years ago (Fig 6D).

**Fig 6.**
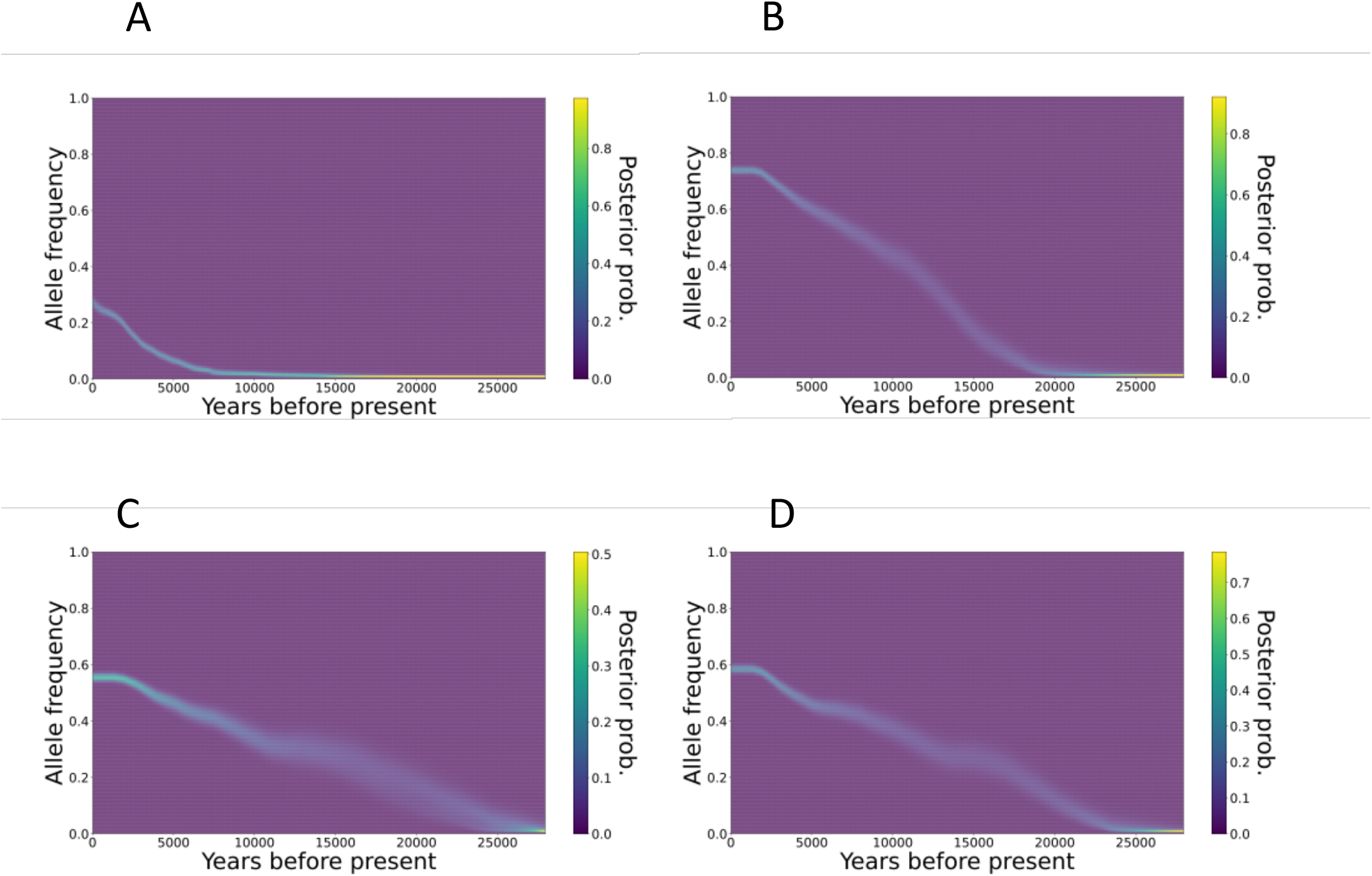
Trajectories of allele frequency of genes. Allele frequency trajectories of (A) ADH1B rs1229984, (B) ALDH2 rs671, (C) OCA2 rs1800414, and (D) FADS1 rs174599 are shown.

## Discussion

In the present study, we conducted a WGS analysis of samples from five biobanks in Japan. Although the data obtained in this study are intended to be provided as control data for genomic studies of various diseases, the analysis in this study focused on data quality and population genetics properties. A uniform quality of data was obtained through the use of a single procedure that encompassed both sequencing and data analysis. Population-based studies using WGS analysis have been conducted in various populations [5,28,29]. Studies on Japanese populations have already been reported [28], and the allele frequency distributions in previous studies are consistent with the results of the present study (Fig 3A). The samples analyzed in the present study were provided by biobanks in three regions of Japan: NCGM, NCCHD, and NCNP in the Tokyo area; NCGG in Aichi Prefecture in the central area in the Honshu Island; and NCVC in Osaka Prefecture in the western area in Honshu Island. Therefore, the genomic information obtained in the present study is expected to reflect the regional diversity of Japan to some extent. For instance, the population genetic analysis identified two clusters representing the ancestry of Ryukyu Islands, comprising Okinawa Prefecture and the islands of Kagoshima Prefecture and the Hondo region (mainland). This supports the idea that the Hondo and Ryukyu populations are genetically differentiated, as suggested by anthropological studies [7–9]. We further found that past population sizes differed between Hondo and Ryukyu. There was a reduction in the Hondo population from 17 to 11 generations ago (Fig 5A). The corresponding period was 476 and 308 years ago, and the assumption is that each generation spanned 28 years; most of this duration overlaps with the Edo period in Japan. This is consistent with findings from historical demography studies, which suggest that the population not only increased but also remained stagnant due to limited economic growth, population concentration in cities, and famine caused by cold weather-related damage during this period. In contrast, the Ryukyu populations showed population reduction from 100 generations ago to 25 generations ago but then increased until the present (Fig 5A).

This population growth about 700 years ago was close to the beginning of farming in the Ryukyu Islands (12th century). Assuming that agriculture was brought to the Ryukyu Islands by migrants from the mainland of Japan, the population decline observed in the Ryukyu population can be considered a bottleneck associated with the migration. The population size estimated from the modern genome reflects the past population of the migrants and should be influenced negligibly by the genetic diversity of the original inhabitants of the Ryukyu Islands. Indeed, although the several human skeletal remains have been discovered from Pleistocene sites in Ryukyu Islands [30, 31], the previous population genetic analysis based on genome-wide SNPs suggested minor genetic contribution of the Pleistocene Ryukyu Island population to the modern Ryukyu population [15, 16]. The estimated time of divergence between Hondo/CHB and Ryukyu was 3,700 years ago (S4 Fig), suggesting that migration to the Ryukyu Islands occurred recently.

Studies of rare genetic diseases require data on the frequency of variants in the population. Most of the variants we found in this study were rare, and many of them were newly discovered in this study, as expected from population genetics theory. However, the lower the frequency of the variants, the more difficult it becomes to distinguish them from errors. In this study, we evaluated the accuracy of genotype detection, estimating a discordance rate of 0.063% compared to genotype detection using WGS and SNP arrays. However, this is an overestimation of the error rate due to the combined error of both technologies. We also used the data obtained from the WGS analysis of the trio for validation. We estimated the Mendelian error rate, which is the proportion of genotypes detected in the offspring of a trio that is inconsistent with Mendel’s laws of heredity. This method has the advantage of being able to examine the entire genome compared to the use of SNP arrays. We found that the Mendelian error rate is much higher for novel variants, i.e., previously reported variants. The Mendelian error rate for novel SNVs was 2.26%, much higher than that of the known SNVs (0.09%). This has important implications for the identification of causative mutations in rare genetic diseases, as many causative mutations for these conditions are newly discovered rare variants. This means that the discovery of such pathological variants in patient sequencing is subject to a non-negligible degree of error.

We conducted the functional annotation of the variants discovered in this study. Consistent with previous studies [5,28,29], variants that were expected to have a high biological impact were less common in the population, confirming that negative natural selection shapes the diversity of variants. Most of the LoF mutations were extremely rare, and most of them were heterozygous (Fig 4A). The number of LoF mutations in the genome was comparable to that in other Eurasian populations (Fig 4B). Although the Ryukyu population has experienced population decline (Fig 5A), the frequency of LoF variants was comparable to that in the Hondo population, and no evidence of differences in the profile of rare functional variants due to the bottleneck effect was noted. The number of LoF sites and homozygous LoF per individual in this study were higher than those detected in a previous study [32]. Among these, the number of stop gained SNVs was consistent with that recorded in the previous study [32], whereas the number of splice site SNVs and frameshift INDELs was higher than that in the previous study [32] (S2 Fig). The number of LoF sites was generally consistent with the number of LoF sites before manual curation in a previous study [33]; thus, it may be possible to remove false-positive homozygous LoFs through manual filtering, as in the previous study [33].

We also examined pathogenic variants that have been reported in the past. Pathogenic variants assessed by an expert panel (4-star status) on ClinVar were found only in one to a few individuals in the population. On the other hand, some variants that were less reviewed were polymorphic with high frequency. These results reinforce the importance of utilizing the frequency of the variants in the population to evaluate their pathogenicity.

Genes that have undergone positive natural selection in the East Asian populations are related to the metabolism. This study supported that the dietary changes in the ancestors seem to have shaped gene frequencies. Candidate regions undergoing positive natural selection were found on a genome-wide scale using genealogy analysis (S5 Fig). ADH1B is involved in metabolizing alcohol to acetaldehyde, and ALDH2 is involved in metabolizing acetaldehyde. Both the non-synonymous A allele of ALDH2 rs671 and the C allele of ADH1B rs1229984 affect the retention of acetaldehyde in the body and cause alcohol flush in Asians [17, 18]. These alleles have been suggested to be associated with Japanese dietary habits and diseases, such as esophageal cancer [34, 35]. Previous studies have hypothesized that positive selection may have acted to maintain acetaldehyde in the blood against parasite infection, which correlates with large-scale rice cultivation [36–39]. We also observed that the increase in the frequency of ADH1B occurred earlier than that of ALDH2, indicating that positive selection began to act at different times for these two genes (Fig 6A and 6B). Based on the geographic distribution of haplotype structures around ADH1B and ALDH2, according to Koganebuchi et al., positive selection on ADH1B rs1229984 started before the beginning of the Jomon period, while positive natural selection on ALDH2 began around 8,000 years ago, in association with the beginning of rice cultivation in China [39]. Our dating by genome-wide genealogy of the Japanese population genome is consistent with the above consideration. Using HapMap data, OCA2 rs1800414 has been shown in previous studies to be the effect of positive natural selection on East Asians [19]. For the OCA2 gene, positive natural selection signals were found in the European population for skin color-related SNPs other than those detected in this study [19]. As natural selection works for light skin color, a previous study mentioned that it enhances vitamin D synthesis capacity in regions with low sunlight [20]. For East Asians as well, positive natural selection may have operated in relation to vitamin D synthesis in regions with low sunlight. However, since the derived allele of rs1800414 is not necessarily more frequent at the high latitudes of East Asia, other possibilities, such as sexual selection, cannot be ruled out at this time [19]. The derived allele of rs1800414 has been shown to be associated with light skin color and tanning ability in Chinese and Japanese populations [20, 21] and is widely observed in modern East Asians [19], suggesting that the derived allele of rs1800414 originated in the common ancestor of East Asians and spread throughout East Asia at very early stages of the East Asian population history. We estimated that the derived allele of OCA2 rs1800414 began to increase in frequency around 25,000 years ago (Fig 6C). Future analyses of older East Asian lineages, such as the ancient genome of the Jomon people, may reveal the original variant of this allele that led to positive natural selection. FADS1 and FADS2 participate in fatty acid metabolism. For example, in the Inuit population, which relies heavily on a marine animal diet, there are positive natural selection signals on SNPs of FADS2 genes, which are responsible for the increase in the concentrations of short-chain fatty acids [40]. Signals of positive natural selection on alleles that promote long-chain fatty acid synthesis have also been identified in African [22], European [25,26,41], and South Asian populations [24]. In particular, studies in European populations have shown that the derived alleles of rs174594 and rs1714546 are associated with increased total cholesterol and LDL cholesterol levels, increased expression of FADS2, and decreased expression of FADS1 [25]. In European populations, increased reliance on plant diets seemed to have resulted in positive natural selection on alleles that promote long-chain fatty acid synthesis pathways of the FADS gene family [23,25,26]. The SNPs in the FADS gene family detected in this study were associated with total cholesterol and LDL cholesterol levels, increased expression of FADS2, and decreased expression of FADS1 (S3 Table), like the SNPs subjected to natural selection in the European population (S3 Table). These results suggest that in Hondo populations, as in Europeans, the dietary change was accompanied by positive natural selection for alleles that promote the long-chain fatty acid synthesis. The frequency of the derived allele of rs174599 in FADS2 began to increase around 25,000 years ago, but the increase was not continuous, and there was a period of stagnation from 15,000 years ago for 5,000 years (Fig 6D). Interestingly, the frequency of this allele varies widely among East Asian populations. The derived allele was major in CHB (64%) and Japanese (63%), whereas it was minor in Dai (Chinese Dai in Xishuangbanna, China) (22%), Han Chinese in South (42%), and Kinh Vietnamese (20%); these data suggest that the positive natural selection of the FADS gene family in East Asians may reflect the association with agriculture and the complex dietary differences among regional populations. Notably, Mathieson and Mathieson (2018) disproved the simple idea that these derived alleles underwent positive natural selection in relation to the introduction of agriculture and speculated that there were complex underlying factors, such as unknown dietary changes [26].

In this study, we demonstrated that the data presented here can be used as a foundation for analysis of human genetics. While this study focused on population genetic characterization of the Japanese population, the data can be used in disease studies, as a resource for genotype imputation in studies of common diseases, and as a control in studies on rare diseases.

## Materials and Methods

### Sample preparation

DNA samples stored in the biobanks of five national centers (National Cerebral and Cardiovascular Center; NCVC, National Center for Geriatrics and Gerontology; NCGG, National Center for Global Health and Medicine; NCGM, National Center of Neurology and Psychiatry; NCNP and National Center for Child Health and Development; NCCHD) were submitted for WGS analysis. Samples derived from healthy individuals or patients with some common diseases were selected as control groups for future disease studies. This study was conducted with approval from the ethics review committee of the NCGM. Informed consent for the analysis of these samples was received from all subjects in each biobank. Approximately 50 μl of DNA at a concentration of 80 ng/μl per sample was aliquoted into 96-well plates and shipped to an outsourced laboratory (TakaraBio, Shiga, Japan) for WGS analysis.

### WGS

To avoid quality fluctuations and batch effects, all samples were analyzed by a single outsourced laboratory. WGS analysis was performed using NovaSeq6000 (Illumina, San Diego, CA, US), and sample preparation was performed using the procedures and reagents recommended by the manufacturer. DNA molecules were sonicated with a protocol targeting an average size of 550 bp. DNA libraries were prepared using the TruSeq DNA PCR-Free HT Library Prep Kit, and index sequences were added for multiplex analysis. The insert size was confirmed by electrophoresis in the range of 400–750 bp before sequencing runs. WGS was performed at 150 bp paired-end and repeated in multiplex until an output of >90 Gbases without duplicated reads was obtained.

### Data analysis

We received the quality controlled FASTQ data from the outsourced laboratory and performed mapping and variant calling in an in-house data analysis pipeline. The mapping and variant calls were performed using the Parabricks v3.1.0 (Nvidia, Santa Clara, CA, US), which provides the capability to perform the analysis recommended by GATK at high speed using a GPU [42]. The GRCh38 was used as the reference sequence. The pipeline used in this study implements algorithms equivalent to those of bwa (v0.7.15) [43] and GATK (v4.1.0). We flagged duplicates from mapped reads but did not perform realignment and base quality score recalibration to reduce the computational time. The mapped data were outputted in BAM format and converted into CRAM format using samtools [44] to reduce the file size. Variant calls were output in gVCF format for joint calling. QC metrics were obtained to evaluate the quality of the analyzed data. The depth and map rate were calculated using GATK’s CollectWgsMetrics tool. These QC metrics were continuously monitored throughout the analysis. The sex chromosomes were analyzed assuming both male and female genders. Variant calls were performed for chromosome X in the diploid (-ploidy 2) model for females and the monoploid (-ploidy 1) model for males. Variant calls of the pseudoautosomal region were performed in the diploid model. Variants of chromosome Y were called in the monoploid mode regardless of the sample’s gender.

Finally, the appropriate gVCF file for each sex was used during joint calling. Data from the high-coverage WGS analysis of 2,504 individuals of the International 1000 Genomes Project phase 3 [10] were used as population references for this study. The CRAM files reanalyzed using high depth WGS were downloaded from a public database, and variant calls were performed with the protocol described in this study.

### Integrated analyses

To properly estimate the frequencies of variants found after WGS in the population, we integrated the gVCF files. The joint calling was conducted by combining samples from the biobank of NCBN and samples from the International 1000 Genomes Project phase 3. For the joint calling, we used the gVCFtyper program of the Sentieon package [45]. This program produces results equivalent to those of GeomicsDBImport followed by GenotypeGVCFs programs for the joint calling of GATK. To perform efficient computation in a cluster computation environment, we divided the autosomes into 29 regions evenly. Each variant was scored using VQSR to filter the integrated VCF. The VarCal and ApplyVarCal programs of the Sentieon package corresponding to GATK’s VariantRecalibrator and ApplyVQSR, respectively, were used for this process. The HapMap and International 1000 Genomes Omni2.5 sites, the high-confidence SNPs of the International 1000 Genomes Project, and the dbSNP151 sites were used as true, training, and known datasets, respectively. Variants identified as PASS, which correspond to filtering with 99.9% sensitivity, were used in subsequent analyses unless otherwise noted. The INDELs in the present variant set were normalized by performing left align, and multiallelic variants were split into multiple variant records using the norm subcommand in bcftools [44].

### Variant annotation

Variants were annotated with the Variant Effect Predictor v102 [46]. We ran the loftee plugin to evaluate the effects of the LoF variants. For the other evaluation of the functional effects, dbNSFP4.1 [47] was used to assign precomputed evaluation values to the variants. The metrics used for the assignment included LRT, SIFT, MutationTaster, and Polyphen2.

### Genotyping by SNP array

Using JaponicaArray [48], genome-wide genotyping was performed on a subset of samples for comparison with WGS results. Ninety-four samples each from five biobanks were analyzed using the residual DNA after WGS analysis. The analysis was performed by an outsourced laboratory (Toshiba, Tokyo, Japan), and the raw data in CEL format was received. Four samples were dropped from the genotyping due to a low call rate (<97%) in the first step of genotyping. Clustering for SNP genotyping of variants was performed on the data of 466 individuals using the Analysis Power Tools (ver. 2.10.2.2, Thermo Fisher Scientific, MA, USA). The clustering results for each probes’ intensity were classified using the SNPolisher program bundled with the Analysis Power Tools, and the 639,508 SNPs classified as “Recommended” in autosomes were used for subsequent analyses. The genotype concordance with WGS was estimated using the hap.py software. To compare the positions for which probes were designed in the SNP array, SNVs with the same position as the SNP array were extracted from the results of WGS analysis. The SNP array results were used as true data and the WGS results as query data. The genotype discordance rate between the SNP array and WGS was calculated by dividing the number of false positives by 639,508, which is the total number of SNPs compared.

### Allele frequency estimation

To calculate the accurate allele frequencies, the ancestry of the samples was estimated using PCA. Variants were filtered under more stringent criteria for this purpose.

Individual’s genotypes were considered no calls if they had a genotype quality (GQ) of less than 20, a depth outside the range of 11 to 64, or if less than 25% of the reads supported the minor allele for heterozygous calls. Then, sites with SNPs that had a VQSR filter of PASS, a minor allele frequency of >1%, and a call rate of >95% were retained. The KING program [49] selected samples of unrelated individuals in the third-degree kinship or more. For this dataset, independent SNPs were extracted using PLINK1.9 [50] with “-indep-pairwise 500 50 0.1”, and PCA was performed to calculate the principal component values for each sample using PLINK1.9 [50]. Clusters were identified visually on the scatter plot of the first and second principal components.

Allele and genotype frequencies were estimated for each ancestry group and biobank. The fill-tags plugin of bcftools was used for these calculations. To compare the allele frequencies in the Japanese population, we downloaded the GEM Japan frequency panel information from TogoVar. Since the GEM Japan panel only provides information in hg19 coordinates, we converted it to GRCh38 coordinates. We used GATK’s LiftoverVcf program for the conversion.

### HLA analysis

Three-field HLA alleles calling was performed using HLA-HD v1.3.0 [51] based on IPD-IMGT/HLA v3.43.0 [52]; a score based on the weighted read counts considering variations in and outside of the domain for antigen presentation was calculated to select the most suitable pair of alleles amongst the candidate HLA alleles. To validate the accuracies of HLA calling from WGS, HLA allele frequency distribution was compared with the HLA frequency dataset from HLA Foundation Laboratory (Kyoto, Japan). To evaluate the accuracy of HLA calling from the WGS dataset, a subset of the samples (n = 94) was subjected to high-resolution experimental HLA genotyping for eight HLA genes (HLA-A, - C, -B, -DRB1, -DQA1, -DQB1, -DPA1, and -DPB1) using next-generation sequencing and AllType assay (One Lambda, West Hills, CA, US). Experimental HLA genotyping was carried out following the vendor instructions, which consist of HLA gene amplification, HLA library preparation, HLA template preparation, and HLA library loading onto an ion 530v1 chip (Thermo Fisher Scientific) in the Ion Chef (Thermo Fisher Scientific), followed by final sequencing on the Ion S5 machine (Thermo Fisher Scientific). HLA genotype assignments were carried out using HLATypeStream Visual (TSV v2.0; One Lambda, West Hills, CA, US) and NGSengine® (v2.18.0.17625, GenDX, Utrecht, the Netherlands).

### Haplotype phasing

Variant phasing was performed using shapeit v4.2 [53] in a haplotype-based analysis. SNPs of unrelated samples identified using the ancestry inference were extracted for phasing. The variant phasing was performed by dividing the autosomes into regions containing overlaps for efficient computation. Each region was about 10 Mb in length, with a 500 kb overlap margin at both ends. After phasing, VCFs were concatenated using the concat subcommand in bcftools.

### Estimation of recent population size change

We estimated the effective population size change of the Japanese population from IBD sharing, which can estimate the population size change in the recent past (∼200 generations ago) using WGS data [13]. Population size change was estimated for each population based on the whole-genome data of Hondo (8,524 individuals) and Ryukyu (182 individuals).

First, the hapibd software [54] was used to detect the IBD segments shared by each individual. For the genetic distance, we referred to the HapMap genetic map data distributed with hapibd. We then estimated the population size change of the Hondo and Ryukyu populations using IBDNe (ibdne.23Apr20.ae9.jar). The shortest threshold of the IBD segment length was set at 2 cM.

### Estimation of genome-wide genealogy, estimation of population size change, and detection of positive natural selection

We conducted the analysis of gene genealogy using the Relate software [14] to detect long-term population size change and positive natural selection in Hondo and Ryukyu populations. Relate is a software that can estimate genealogy on a genome-wide scale for over 10,000 samples [55]. In this study, we used 1,000 randomly selected individual genomes of Hondo, 182 Ryukyu samples, and 103 CHB samples of 1000 Genomes Project [10]. First, input files (.haps, .samples) were created from vcf files using the PrepareInputFiles.sh script in Relate software. We retrieved the Homo sapiens ancestral sequences (GRCh38) of Ensembl 103 for the ancestral sequence and StrictMask of 1000 Genomes Project for genomic mask. Next, genome-wide genealogy was estimated using the “Relate” command of Relate software packages. The mutation rate was set to 1.25 × 10^- 8^ per base per generation and the effective population size was set to 30,000. We assumed 28 years as the generation time in humans. The estimated genome-wide genealogy (.anc, .mut) was used as input for population size estimation of Hondo and Ryukyu populations using the EstimatePopulationSize.sh script. This script simultaneously conducts estimation of population sizes, re-estimation of branch lengths using the estimated population sizes, and estimation an average mutation rate. Finally, based on genome-wide genealogy, we detected the target SNPs of positive natural selection acting on Hondo and Ryukyu populations. Relate calculates a p-value of each SNP for positive selection that quantifies how quickly a mutation has spread in the population. The p-values were calculated for each SNP using the DetectSelection.sh script using the output genealogies of population size estimation (.anc, .mut). We evaluated the quality of each SNP by “RelateSelection –mode Quality,” and SNPs inferred to be inaccurate tree estimation were excluded.

### Estimating the allele frequency trajectory

Changes in the allele frequency through the time were estimated using CLUES to infer allele frequency trajectories [27]. CLUES uses the genome-wide genealogy inferred by Relate. First, the sampleBranchLengths.sh script implemented in Relate was used to MCMC sample the gene trees for the focal SNPs. Then, using the sampled tree file (.timeb) as input, we estimated the allele frequency trajectory using CLUES’s inference.py command. The coalescence rate estimated by Relate (.coal file) can be used as an input to modify the population size change using the -coal option. In this study, we estimated the allele frequency trajectory by focusing on ALDH2 rs671, ADH1B rs1229984, OCA2 rs1800414, and FADS2 rs174600 among the SNPs that showed signals of natural selection in Relate.

## Acknowledgments

We thank the participants who provided biological samples for this study. We thank Ms. Megumi Tatsumi, Ms. Haruna Kaneko, Ms. Ayumi Fujisawa, Ms. Mami Sasaki, Ms. Yukiko Miyashita, and Ms. Chikako Kinjyo at NCNP Biobank; Ms. Sachiyo Ito at NCGG Biobank; Dr. Satoshi Suzuki at NCGM Biobank; the personnel at NCVC Biobank; Ms. Kazuyo Kawamura, Dr. Kumiko Yanagi, Dr. Saki Aoto, and Ms. Fuyuki Hasegawa at NCCHD Biobank; and Ms. Akino Takase at NCBN Central Biobank for their contributions to this study. We also thank Dr. Leo Speidel for discussion on the results of Relate analysis and Dr. Osamu Ogasawara for the technical support for the development of secondary analysis pipelines. Computations were partially performed on the NIG supercomputer at ROIS National Institute of Genetics.

## Author contributions

**Conceptualization:** Y.K., N.E., M.M. and T.K.

**Methodology:** Y.O., R.Mi., E.N., H.G., K.Hata., K.Hatt., A.I., H.I-U., T.Kana., T.Kant., R.Ma., M.N., K.O., M.S., A.T., H.T., T.T., A.U., H.W., S.Y., Y.G., Y.Mar., Y.Mat., and S.N.

**Data Curation:** K.K., H.S. and K.M.

**Formal Analysis:** Y.K., Y.W., and S-S.K.

**Project Administration:** Y.G., Y.Mar., Y.Mat., S.N. and K.T.

**Writing – Original Draft Preparation:** Y.K., W.Y., Y.O. and T.K. Y.K., N.E., M.M., and T.K. designed the study. Y.O., R.Mi., H.G., K.Hata., K.Hatt., A.I., H.I-U., T.Kana., T.Kant., R.Ma., M.N., K.O., M.S., A.T., H.T., T.T., A.U., H.W., S.Y., Y.G., Y.Mar., Y.Mat., and S.N. contributed to the whole-genome sequencing. Y.O., R.Mi., and E.N. contributed to the SNP genotyping. K.K., H.S., and K.M. contributed to the data collection. Y.K., Y.W., and S-S.K. contributed to the data analysis. Y.G., Y.Mar., Y.Mat., S.N., and K.T. contributed to the management of biobank. Y.K., W.Y., Y.O., and T.K. wrote the manuscript with input from all authors.

## Supporting information

Supporting information includes seven figures and three tables.

## Web resources

IGSR: The International Genome Sample Resource, https://www.internationalgenome.org Hap.py, https://github.com/Illumina/hap.py

GEM Japan, https://www.amed.go.jp/en/aboutus/collaboration/ga4gh_gem_japan.html TogoVar, https://togovar.biosciencedbc.jp/

GATK, https://gatk.broadinstitute.org/hc/en-us

HLA frequency dataset from HLA Foundation Laboratory (Kyoto, Japan), http://hla.or.jp/index.html

Homo sapiens ancestral sequences (GRCh38), ftp://ftp.ensembl.org/pub/current_fasta/ancestral_alleles/homo_sapiens_ancestor_GRCh38.t ar.gz

dbSNP, https://www.ncbi.nlm.nih.gov/snp/

## Data and code availability

The allele and genotype frequency data are available in the NBDC human database; Accession: hum0331. The raw genomic data are available upon request to corresponding authors and will soon be shared on a computational infrastructure currently under construction by the Japan Agency for Medical Research and Development.

## Supporting information

**S1 Fig. Genetic structure of East Asian populations.** The clusters consisting of the NCBN samples in Figure 2 are classified into Hondo (black), Ryukyu (orange), and others (blue).

**S2 Fig. Analysis of Loss-of-function (LoF) variants.** The numbers of LoF sites per individual by category are presented: (A) and (B) stop gained SNV; (C) and (D) splice site SNV; (E) and (F) frameshift INDELs.

**S3 Fig. HLA alleles frequencies (%) between NCBN vs HLA Foundation Laboratory, Kyoto, Japan.** Comparison for class I HLA genes (HLA-A, -C, -B) (left). Comparison for class II HLA genes (HLA-DRB1, -DQA1, -DQB1, -DPA1, -DPB1) (right). Only common HLA alleles (HLA frequencies > 1%) are included in this analysis.

**S4 Fig. Long-term effective population size change of Hondo, Ryukyu, and Han Chinese.** The changes in population size were estimated from the gene genealogy across the genome.

**S5 Fig. Manhattan plot of the selection scan result of the whole-genome SNPs by Relate.** The red line represents the genome-wide significance level (5 × 10^-8^).

**S6 Fig. QQ plot of the selection scan result of the whole-genome SNPs by Relate.** The red line denotes y=x.

**S7 Fig. Gene genealogy estimated by Relate.** Genealogy of (a) ALDH2 rs671, (b) ADH1B rs1229984 (c) OCA2 rs1800414 (d) FADS1 rs174599 are presented. The vertical axis represents the age (years before present). Derived allele carriers are shown in red.

