## Supplemental Materials for "Exploring the genetic diversity of the Japanese Population: Insights from a Large-Scale Whole Genome Sequencing Analysis"

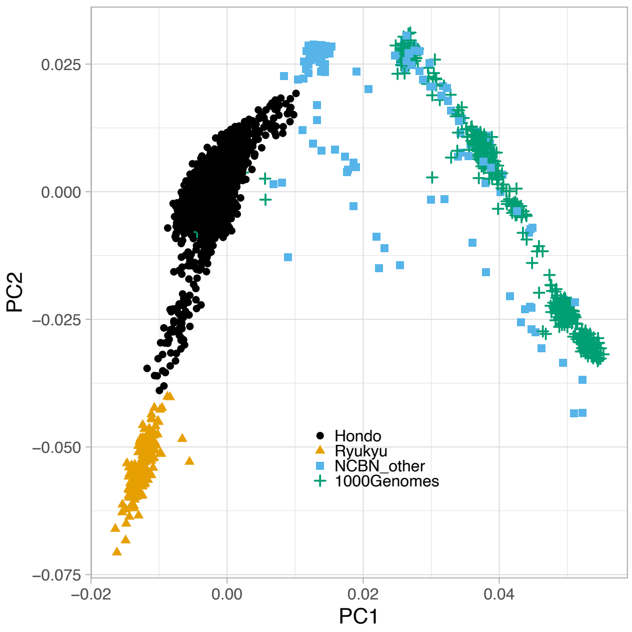


Figure S1: **Genetic structure of East Asian populations**
The clusters consisting of the NCBN samples in Figure 2 are classified into Hondo (black), Ryukyu (orange), and others (blue).

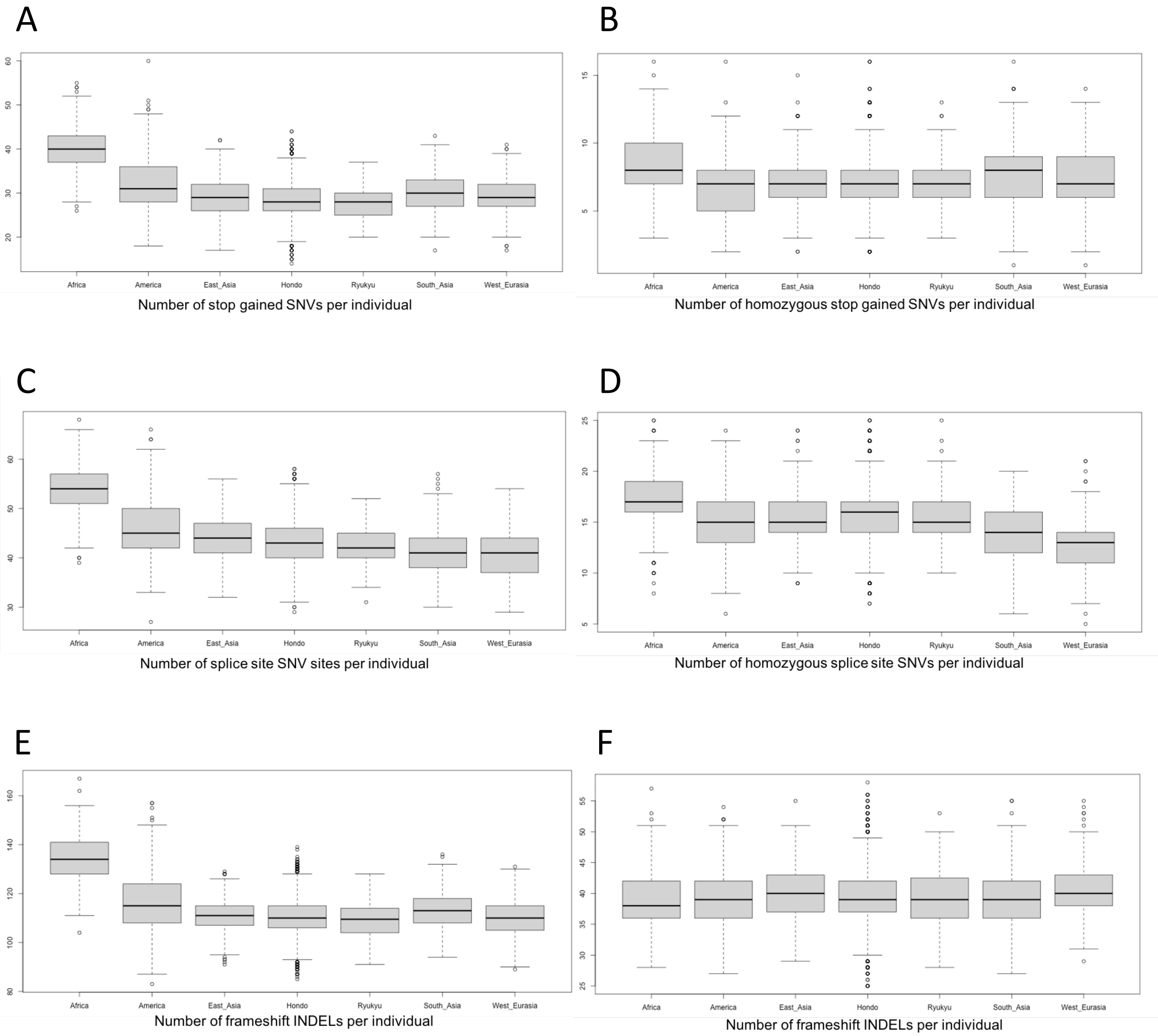


Figure S2: **Analysis of Loss-of-function (LoF) variants**.
The numbers of LoF sites per individual by category are presented: (A) and (B) stop gained SNV; (C) and (D) splice site SNV; (E) and (F) frameshift INDELs.

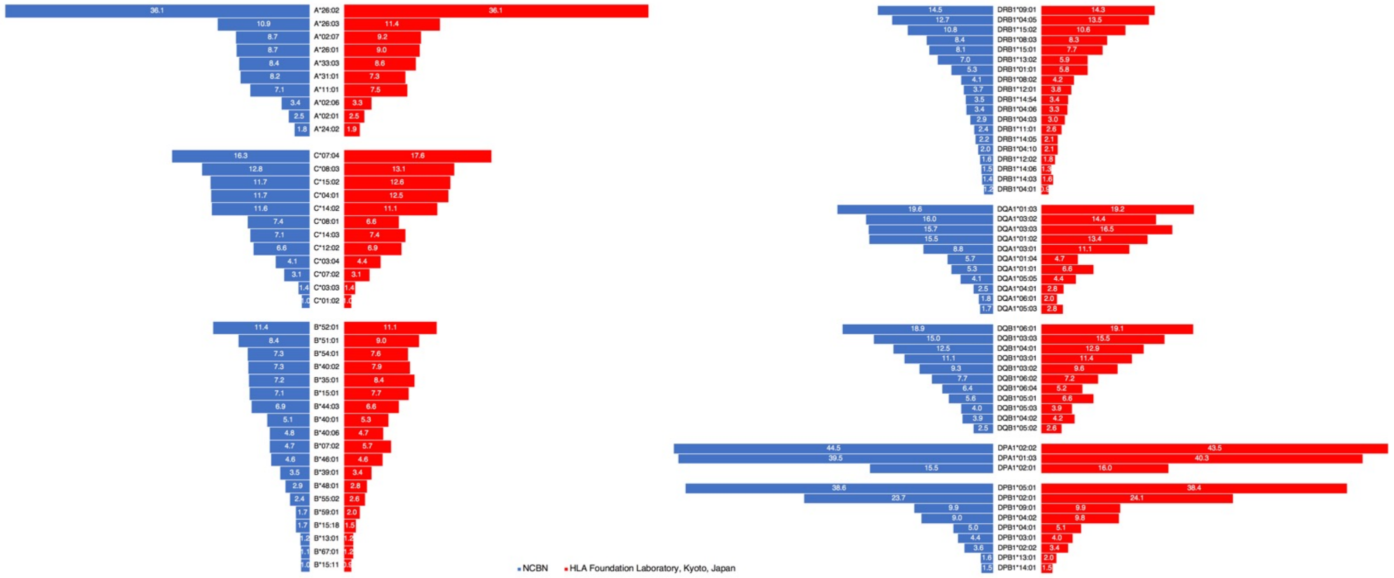


Figure S3: **HLA alleles frequencies (%) between NCBN vs HLA Foundation Laboratory, Kyoto, Japan**.
Comparison for class I HLA genes (HLA-A, -C, -B) (left). Comparison for class II HLA genes (HLA-DRB1, -DQA1, -DQB1, -DPA1, -DPB1)(right). Only common HLA alleles (HLA frequencies > 1%) are included in this analysis.

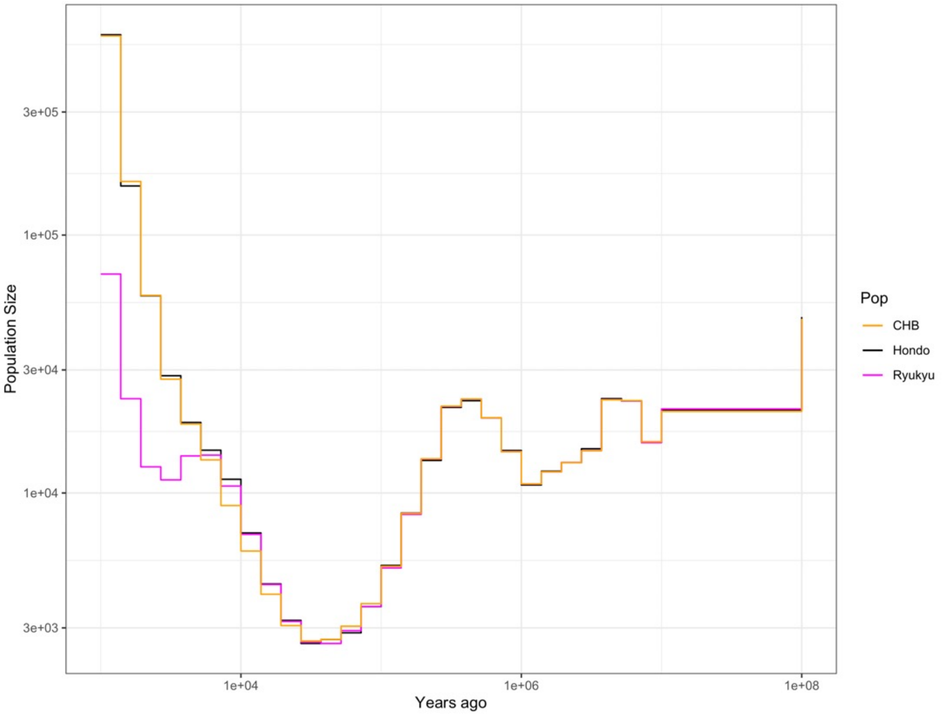


Figure S4: **Long-term effective population size change of Hondo, Ryukyu and Han Chinese.**
The changes in population size were estimated from the gene genealogy across the genome.

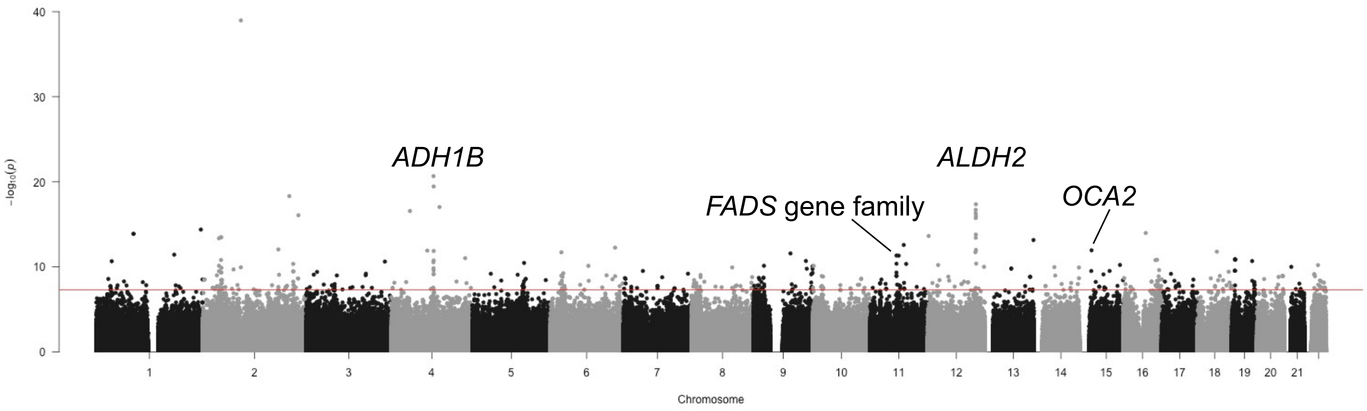


Figure S5: **Manhattan plot of the selection scan result of the whole genome SNPs by Relate.**
The red line represents the genome-wide significance level ($5 \times 10^{-8}$).

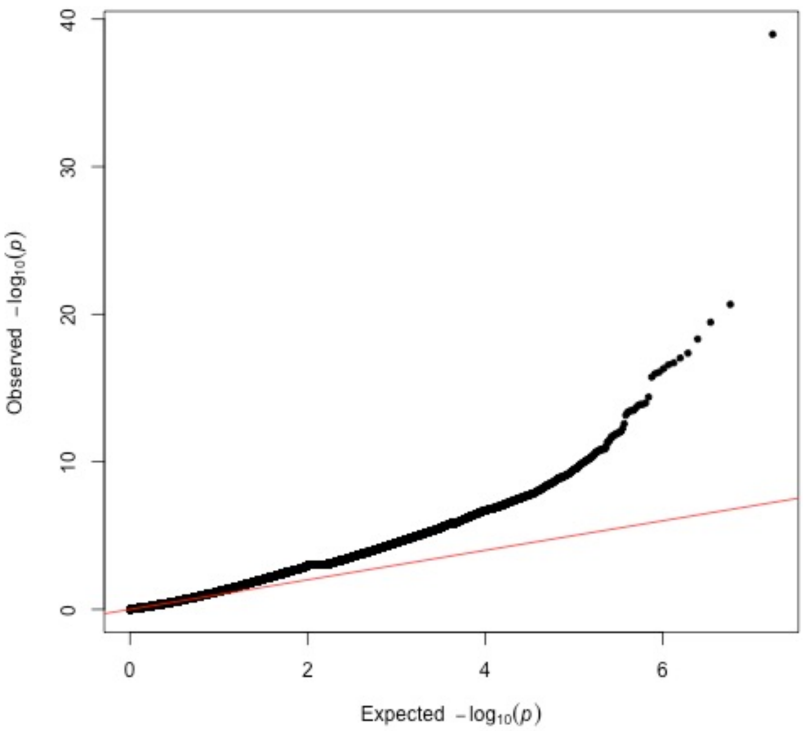


Figure S6: **QQ plot of the selection scan result of the whole genome SNPs by Relate.**
The red line denotes y=x.

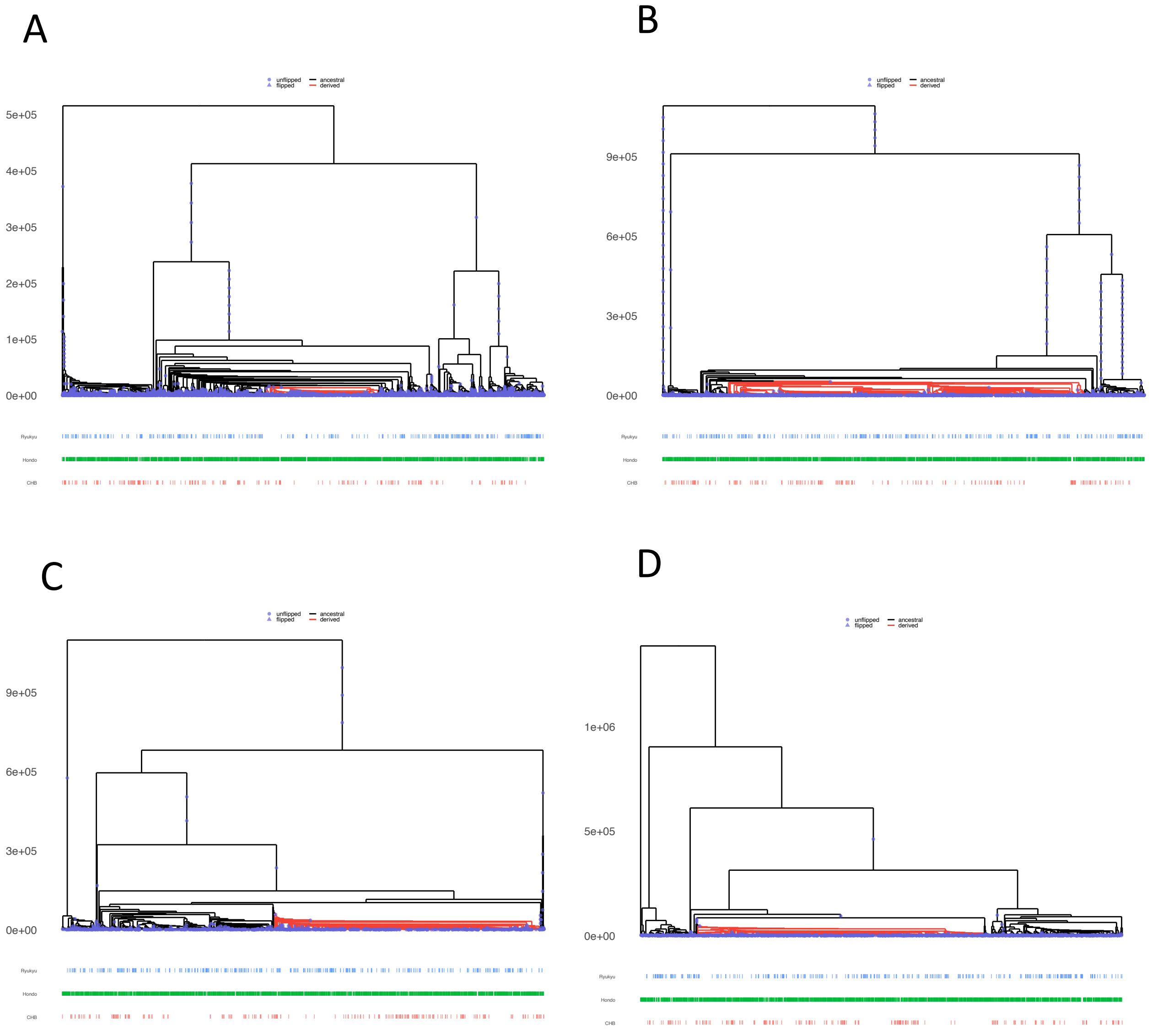


Figure S7: **Gene genealogy estimated by RELATE.** Genealogy of (a) ALDH2 rs671, (b) ADH1B rs1229984 (c) OCA2 rs1800414 (d) FADS1 rs174599 are presented. The vertical axis represents the age (years before present). Derived allele carriers are shown in red.

| Data set | typer of variant | novelty (dbSNP151) | filter(VQSR) | number | $F_{singleton}$ | $F_{MAF<0.5\%}$ |
|---|---|---|---|---|---|---|
| NCBN+1000G | SNV | known | PASS | 106,565,530 | 41.11% | 93.77% |
| | SNV | known | FAIL | 13,086,683 | 26.93% | 92.96% |
| | SNV | novel | PASS | 50,408,517 | 71.92% | 99.99% |
| | SNV | novel | FAIL | 10,197,934 | 40.16% | 99.52% |
| | INDEL | known | PASS | 17,214,879 | 20.19% | 87.42% |
| | INDEL | known | FAIL | 193,890 | 15.63% | 89.05% |
| | INDEL | novel | PASS | 10,675,637 | 53.53% | 99.44% |
| | INDEL | novel | FAIL | 442,789 | 34.25% | 98.84% |
| | | Total | | 208,785,859 | 46.49% | 95.28% |
| NCBN | SNV | known | PASS | 46,199,619 | 34.56% | 86.73% |
| | SNV | known | FAIL | 7,977,505 | 23.90% | 89.79% |
| | SNV | novel | PASS | 41,046,547 | 67.46% | 99.99% |
| | SNV | novel | FAIL | 8,268,223 | 36.32% | 99.49% |
| | INDEL | known | PASS | 11,122,104 | 19.71% | 82.60% |
| | INDEL | known | FAIL | 149,334 | 11.12% | 87.44% |
| | INDEL | novel | PASS | 7,361,318 | 47.97% | 99.34% |
| | INDEL | novel | FAIL | 334,657 | 31.30% | 98.77% |
| | | Total | | 122,459,307 | 44.43% | 92.65% |

Table S1. **Summary of variants discovered by WGS.**

Provided on a separate EXCEL file (TableS2.xlslx)

Table S2. **List of SNPs for which natural selection was detected by Relate**

| rs ID | Gene | P value | Effect size | Tissue |
|---|---|---|---|---|
| rs174599 | FADS2 | 3.90E-41 | 0.6 | Whole Blood |
| rs174599 | FADS2 | 1.20E-24 | 0.44 | Esophagus - Muscularis |
| rs174599 | FADS1 | 2.60E-24 | -0.36 | Esophagus - Mucosa |
| rs174599 | FADS1 | 5.00E-24 | -0.71 | Brain - Cerebellum |
| rs174599 | FADS1 | 1.50E-22 | -0.56 | Pancreas |
| rs174600 | FADS2 | 1.50E-43 | 0.65 | Whole Blood |
| rs174600 | FADS2 | 7.10E-25 | 0.29 | Cells - Cultured fibroblasts |
| rs174600 | FADS2 | 3.30E-24 | 0.47 | Esophagus - Muscularis |
| rs174601 | FADS2 | 4.40E-41 | 0.6 | Whole Blood |
| rs174601 | FADS1 | 8.50E-25 | -0.36 | Esophagus - Mucosa |
| rs174601 | FADS2 | 2.70E-24 | 0.44 | Esophagus - Muscularis |
| rs174601 | FADS1 | 5.90E-23 | -0.7 | Brain - Cerebellum |
| rs174601 | FADS1 | 1.70E-22 | -0.56 | Pancreas |
| rs97384 | FADS2 | 4.50E-42 | -0.62 | Whole Blood |
| rs97384 | FADS1 | 6.50E-25 | 0.62 | Pancreas |
| rs97384 | FADS1 | 4.80E-22 | 0.68 | Brain - Cerebellum |
| rs97384 | FADS1 | 2.40E-21 | 0.35 | Esophagus - Mucosa |
| rs97384 | FADS2 | 3.10E-21 | -0.42 | Esophagus - Muscularis |

Table S3. Abbreviated GTEx eQTL Results of SNPs affected by positive natural selection in FADS gene family, P Value Cut Off of $10^{-20}$
